# Adaptive algorithms for shaping behavior

**DOI:** 10.1101/2023.12.03.569774

**Authors:** William L. Tong, Anisha Iyer, Venkatesh N. Murthy, Gautam Reddy

**Affiliations:** School of Engineering and Applied Sciences, Harvard University, Cambridge, MA, USA; University of California Berkeley, CA, USA; Department of Molecular and Cellular Biology, Harvard University, Cambridge, MA, USA; Center for Brain Science, Harvard University, Cambridge, MA, USA; Physics & Informatics Laboratories, NTT Research, Inc., Sunnyvale, CA, USA

## Abstract

Dogs and laboratory mice are commonly trained to perform complex tasks by guiding them through a curriculum of simpler tasks (‘shaping’). What are the principles behind effective shaping strategies? Here, we propose a machine learning framework for shaping animal behavior, where an autonomous teacher agent decides its student’s task based on the student’s transcript of successes and failures on previously assigned tasks. Using autonomous teachers that plan a curriculum in a common sequence learning task, we show that near-optimal shaping algorithms adaptively alternate between simpler and harder tasks to carefully balance reinforcement and extinction. Based on this intuition, we derive an adaptive shaping heuristic with minimal parameters, which we show is near-optimal on the sequence learning task and robustly trains deep reinforcement learning agents on navigation tasks that involve sparse, delayed rewards. Extensions to continuous curricula are explored. Our work provides a starting point towards a general computational framework for shaping animal behavior.

## I. INTRODUCTION

Animal trainers “shape” an animal’s behavior towards a specific sequence of actions [1–4], for example, training a dog to roll, fetch and sit. An untrained animal is unlikely to execute this sequence in the right order, even if it can perform each action separately. One intuitive teaching strategy is to first reinforce the animal for rolling. Once the animal rolls consistently, rolling is no longer reinforced (or becomes variable) and the animal is instead reinforced for successfully fetching after a roll. This iterative process is repeated until the animal learns the right sequence. In some cases, the trainer further breaks down the task or “lures” the animal to carry out the desired action.

Here, a shaping process is essential as the animal will rarely execute the right sequence during innate behavior. This simple intuition highlights a fundamental constraint: learning a particular behavioral sequence through random, unguided exploration is inefficient when the dimensionality of behavior is large, regardless of the learning rule the animal employs. Shaping tackles this issue by iteratively approximating longer bits of the sequence, limiting the search space at every stage of training.

Laboratory animals solving a perceptual discrimination task spend a significant fraction of their training time learning the rules of the task. For example, a free-moving two-action-forced-choice paradigm often involves an animal triggering a stimulus through a nose poke at a particular location, which leads to reward delivery at two possible distal locations in the arena. The spatiotemporal relationship between the nose poke and reward, that paying attention to the stimulus matters for obtaining reward and that there is a temporal cost for a wrong choice are non-trivial rules of the environment that the animal has to learn before learning the perceptual features that distinguish the stimulus sets. Significant attention has been paid to active learning [5–8], which addresses the latter problem of choosing a perceptual stimulus set to efficiently teach the stimulus-outcome relationship. Behavioral shaping, on the other hand, is used to teach the rules of the task and closely reflects the curriculum design process used in education.

A shaping protocol typically involves hand-designing a series of simpler tasks leading to the full task during training. The animal is rewarded for successfully completing an assigned sub-task, and the curriculum progresses once the animal is sufficiently good at completing this sub-task [9–13]. However, it is unclear whether such heuristics are close to optimal even in simple scenarios, or when these strategies might fail. Understanding the principles that drive effective shaping, coupled with closed-loop training strategies, could considerably reduce the training time for both laboratory animals and artificial agents, while providing insight into factors that contribute to slow or fast learning [14, 15]. Our goal is to develop a general computational framework for shaping animal behavior, paying particular attention to the constraints that trainers face.

In machine learning, the importance of shaping-inspired approaches for training agents was recognized early on [16–22]. More recently, numerous *automatic* curriculum learning (ACL) techniques have been developed for training deep reinforcement learning (RL) agents (reviewed in [23]). Within the ACL framework, an autonomous teacher agent determines the distribution of the student’s tasks based on the student’s past behavior. However, these approaches rely on arbitrary control over the agent’s states [24–26], exploration [27–34] or the reward structure [35–37]. For example, a wellknown strategy known as potential-based reward shaping [35] modifies the reward function to expedite learning while preserving the optimal policy. Such a procedure is infeasible in experimental situations where the animal has to interrupt its behavior in order to acquire reward. In other cases, these methods assume that the agent’s performance can be measured on a range of arbitrary test tasks [38] or require access to expert demonstrations [26, 39, 40].

Although these assumptions are reasonable for training artificial RL agents and have demonstrated success in numerous tasks, they are not suitable for training animals. When training animals, we typically have 1) limited flexibility in controlling rewards and exploration statistics, 2) partial observability, as animals can often be evaluated based only on whether they have succeeded or failed on the task (their true “state” remains unknown), and 3) no delineation between training and test trials. In addition, animals often have an innate repertoire of responses and behaviors they may resort to by default, and a training procedure which recognizes and takes advantage of this feature may be more successful.

## II. FRAMEWORK

To address these issues, we propose an ACL framework, which we term *outcome-based curriculum learning* (OCL). In OCL, a teacher agent decides the student’s next task based solely on the student’s outcomes, i.e., its history of successes or failures on past tasks, with the long-term goal of minimizing the time to reach a desired level of performance on the final task. By observing and delivering rewards based on binary outcomes, teacher algorithms are task-agnostic and can be applied for training both artificial agents and animals. Closest to our framework is the teacher-student curriculum learning framework [41, 42], which relies on observed scores. Inspired by the concept of learning progress [43] in developmental psychology, Matiisen et al [41] propose heuristic strategies where the teacher selects the task on which the student shows the greatest improvement on scores. However, we find below that these heuristics perform poorly compared to our simpler alternatives.

To gain intuition, it is helpful to visualize teaching with OCL as ‘navigation’ through an (unknown) difficulty landscape that is shaped by the student’s innate biases towards performing behaviors pertinent to the task. Such a difficulty landscape is illustrated in Figure 1a for a task whose difficulty increases along two independent axes. We define difficulty as the negative log probability of success on a task (here parameterized by the two skill axes) *given* the student’s current policy. The difficulty landscape thus depends on the task as well as the student’s innate biases and learned behavior. The goal of OCL is to progressively flatten regions of the landscape to solve the full task as quickly as possible.

**FIG. 1:**
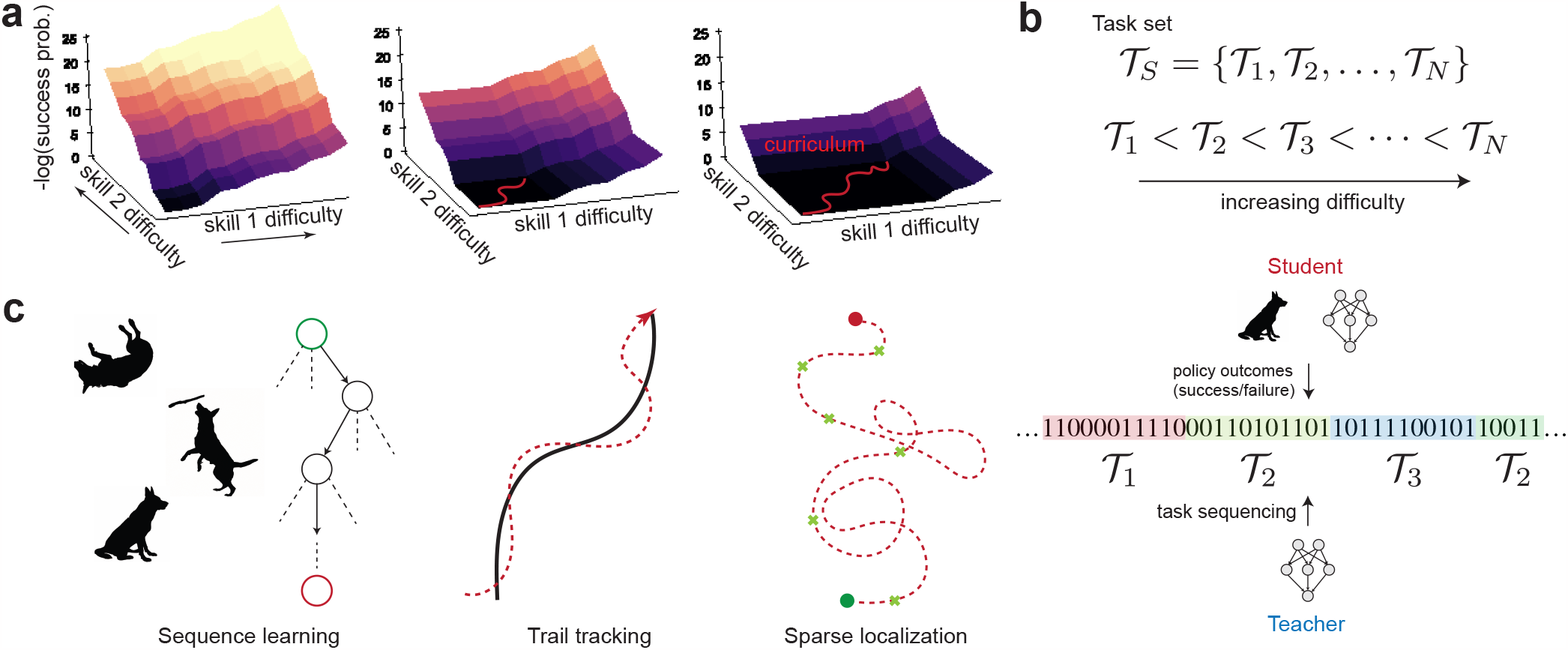
(a) Teaching using our OCL framework can be visualized using a difficulty landscape (here, parameterized by two skill axes), which quantifies the student’s success probability for each difficulty level. A student assigned an extremely difficult task will not learn, since they are unlikely to succeed and thus do not receive significant reinforcement. The teacher’s purpose is to adaptively assign tasks (shown in red) while simultaneously inferring the difficulty landscape to flatten it as quickly as possible. (b) Tasks from a pre-defined set are ordered based on their difficulty, as measured by the success probability of a naive agent. An autonomous teacher decides the student’s task (𝒯_1_, *𝒯*_2_, *𝒯*_3_ … ) based on the student’s transcript of successes and failures (represented here as 0s and 1s respectively) on previously assigned tasks. (c) We apply our OCL framework to three biologically relevant goal-oriented tasks involving delayed rewards: a generic sequence learning task, an odor-guided trail tracking task and a plume-tracking task involving localization to a target based on sparse cues.

In this manuscript, we consider tasks that can be decomposed into a single difficulty scale. Such tasks lend themselves naturally to a curriculum. A student begins with the simplest version of the task and progresses through difficulty levels (as set by the curriculum) until they succeed at the entire task. In the discrete version of OCL, the experimenter designs tasks and rates them based on their difficulty in discrete levels from 1 to *N* (Figure 1b). A desired threshold level of performance is specified for the *N* th task (the full task). Given this input, the teacher algorithms that we consider below choose the appropriate difficulty level for the student based on their past transcripts. At the start of every interaction, the teacher receives as input a transcript and proposes the difficulty level *k*. The student attempts the task for *T* (fixed) rounds, adding to the performance transcript. This two-way interaction continues until the student attains a satisfactory level of success on the final task.

We first investigate in detail a sequence learning task, where an RL-based student is required to learn the correct sequence of *N* actions (Figure 1c). The sequence learning task encompasses a large variety of behavioral tasks, including tricks such as the roll→fetch→sit sequence described above, numerous Skinnerian tasks, as well as common laboratory behavioral experiments which have a self-initiated trial structure. The difficulty landscape of such tasks is determined by the complexity of the sequence (*N* ) and the innate probability that the student will execute the correct action at each step of the sequence. Since the probability of success decreases exponentially with *N*, the agent is unlikely to learn the full task without shaping when *N* is sufficiently large.

The simplicity of the task structure allows us to examine normative teacher strategies using modified Monte Carlo planning algorithms for decision-making under uncertainty. Using insights from these normative strategies, we use differential evolution to design near-optimal heuristics that are agnostic to the task, learning rule, and learning parameters. Next, we apply our method on two novel, naturalistic, sequential decision-making tasks that involve delayed rewards: odor-guided trail tracking and plume-based odor localization (Figure 1c). We show that deep reinforcement learning agents can be trained using our adaptive teacher algorithms to solve these tasks using only a single reward delivered at the end of the task. Finally, we extend this framework to continuous parameterizations of the task, where the teacher has the option of breaking down the task into simpler components.

## III. RESULTS

### A. Sequence learning

In the sequence learning task, a student RL agent begins each trial at a fixed start state and receives a reward *r* if they perform the correct sequence of *N* actions (Figure 2a, see Appendix A for full details). If the student fails to take the correct action at any step in the sequence, the student receives no reward and the episode terminates. The probability that the student takes the correct action at step *i* is given by *σ*(*q*_*i*_ + *ε*_*i*_), where *ε*_*i*_ is the (fixed) innate bias that determines the probability the student will take the correct action before any learning occurs. *σ* is the logistic function. For example, if the agent prior to learning takes *K* possible actions at step *i* with equal probability and only one of them is correct, we have *ε*_*i*_ =−log(*K*−1). The action value *q*_*i*_ is initially set to zero and updated using a standard temporal-difference (TD) learning rule with learning rate *α*.

**FIG. 2:**
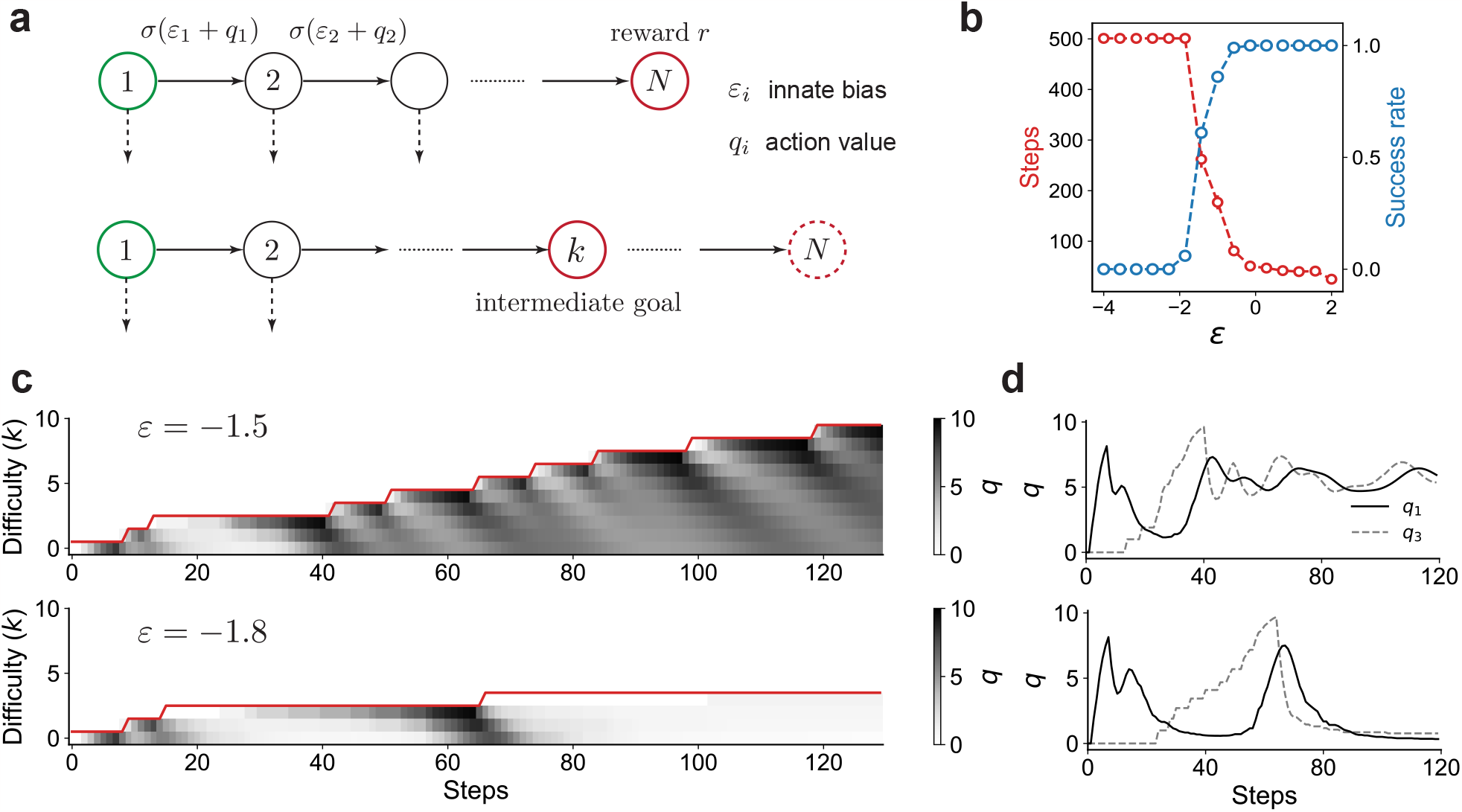
(a) The sequence learning setup. In the full task, the student is required to take a sequence of *N* correct actions to get reward. In intermediate levels of the task, the reward is delivered if the student takes *n ≤ N* correct actions. *ε*_*i*_ is the innate bias of the student to take the correct action at the *i*th step, prior to training. We assume *ε*_*i*_ = *ε* for all *i* unless otherwise specified. (b) The incremental teacher (INC) fails once *ε* ≲*−*1.7. (c) The *q* values (in grayscale) for the correct action at each step shown for *ε* =*−*1.5 (top) and *ε* =*−*1.8 (bottom). The red line shows the assigned task level. Note the striped dynamics in the top row caused due to alternating reinforcement and extinction. In the bottom row, *ε* is too small, forcing learning to stall. (d) Time series of *q* values for actions at the first (solid black) and third (dashed gray) steps for the two examples shown in panel (c).

The sequence learning task naturally splits into discrete difficulty levels: the teacher modulates difficulty by increasing, decreasing or maintaining the step *k* at which the student is rewarded. The innate biases *ε*_*i*_’s play a key role in the dynamics since they determine the probability of success (and thus the rate of reinforcement) when the difficulty level is increased. We assume for simplicity that all *ε*_*i*_’s are equal to *ε*; the general case is considered later. We seek OCL algorithms that minimize the time the student takes to succeed at a rate greater than a threshold *τ* on the full task without prior knowledge of the student’s innate biases and learning parameters.

### B. An incremental teacher strategy is not robust

An intuitive baseline strategy when designing a curriculum is an incremental (INC) approach: the teacher increments the difficulty by one when the student’s estimated success rate *ŝ*exceeds *τ* at the current level. Note that since the success rate changes due to learning, a reasonable estimator *ŝ*should consider recent transcripts yet a sufficient number of them to minimize sampling noise. We consider different estimation procedures for computing *ŝ*and find that an exponential moving average estimator is computationally inexpensive and achieves comparable performance as other more sophisticated methods (Appendix E, Figure S2).

INC is stable for large *ε* (Figure 2b). However, INC abruptly and consistently fails when *ε* is below a threshold (*ε* ≲−1.7 in Figure 2b). Examining the dynamics of the *q* values provides insight into why this catastrophic failure occurs.

Let us first examine *q* value dynamics when the student is required to directly solve the case *k* = 5, where *ε* is chosen such that the student is capable of learning without a curriculum. The dynamics of *q* values exhibit a ‘reinforcement wave’, where actions are sequentially reinforced backwards from the final state to the start [44]. This backward propagation is a generic feature of RL, since the goal acts as the sole source of reward and reinforcement propagates through RL rules that act locally. Now, suppose the difficulty is incremented by one (*k* = 6). Immediately after this change, the student executes the correct sequence of actions until the fifth step, but will likely fail to receive reward as the final step has not been reinforced. These (possibly brief) series of failures produce a long-lasting extinction wave that propagates backwards to earlier steps with dynamics that parallel those of the reinforcement wave. In short, transient failures after every difficulty increment have long-term effects on learning dynamics and success rate.

When visualized over the course of a curriculum, *q* values assume characteristic “striped”-dynamics that emerge due to alternating waves of extinction and reinforcement (top panel in Figure 2c). These striped dynamics reflect the transient failures and eventual successes that follow an increment to higher difficulty when *ε* is larger than the failure threshold. Extinction dominates reinforcement when *ε* is below a critical value, leading to catastrophic unlearning of previous actions and subsequent lack of learning progress. Since extinction is unavoidable after significant increases in difficulty (*e*^*ε*^≪1), optimal strategies that are robust in this regime will have to ameliorate this effect while completing the curriculum as quickly as possible. That is, effective curriculum design strategies should achieve an optimal balance between extinction and reinforcement.

### C . Near-optimal teacher algorithms alternate between difficulty levels

To gain insight into near-optimal strategies, we formulate the teacher’s task for the sequence learning task as optimal decision-making under uncertainty using the framework of Partially Observable Markov Decision Processes (POMDPs) [45–47] . Specifically, the teacher decides whether to increase, decrease or keep the same difficulty level based on the student’s past history, and receives a unit reward when the student crosses the threshold success rate on the full task. A discount factor incentivizes the teacher to minimize the time to reach this goal. As when training animals, one challenge is that the student’s true learning state (encoded by the *q* values) are hidden as the teacher receives only a finite transcript of successes and failures on previously assigned tasks. Another challenge is that the teacher is not *a priori* aware of the student’s innate biases and learning rate. Moreover, the long horizon and sparse reward makes planning computationally prohibitive.

To solve this task, we employ an online POMDP solver (called POMCP [48]) that relies on Monte Carlo planning and inference (Figure 3a). This solver plans based on the inferred joint distribution of *q*’s, *ε* and *α*, which is represented as a collection of particles with different parameter values. A planning algorithm based on Monte Carlo Tree Search (MCTS) [49] balances exploration and exploitation to decide the next action. The student’s transcript on the following round is then used to update the joint distribution using Bayes’ rule implemented as a particle filter, after which this cycle is repeated. With sufficient sampling of particles and planning paths, the solver provides a near-optimal adaptive teacher algorithm for the sequence learning task. Due to the large size of our POMDP, the implementation of POMCP is nontrivial, with details in Appendix E and F.

**FIG. 3:**
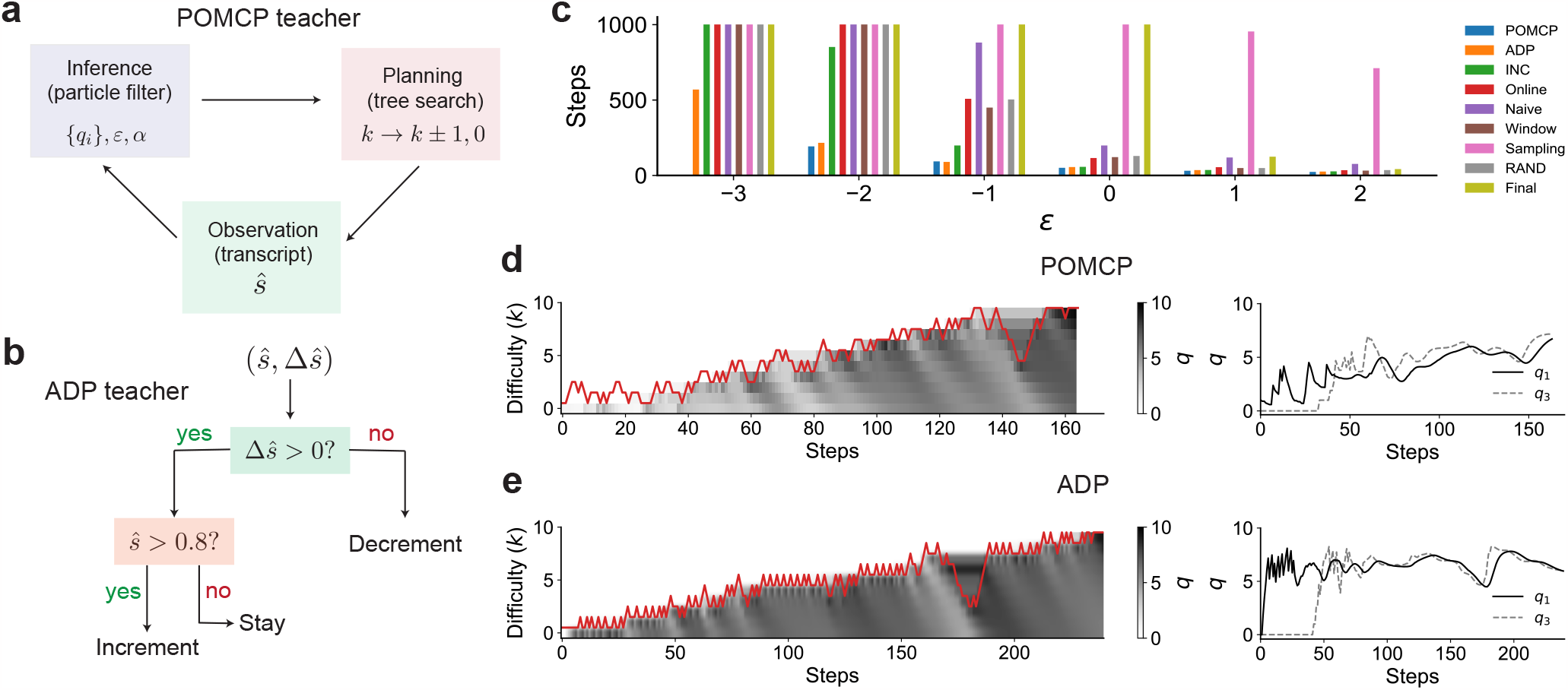
(a) An overview of the POMCP teacher, which cycles between inferring the student’s *q* values, innate bias and learning rate based on the transcript and planning using a Monte Carlo tree search. (b) The adaptive heuristic (ADP), which employs a simple decision rule to stay, increment or decrement the current difficulty based on the estimated success rate *ŝ*(computed using an exponential moving average over past transcripts). (c) POMCP and ADP are comparable and significantly outperform other algorithms [41] when the task is non-trivial (low *ε*), including when INC fails (*ε* ≲ *−*1.7). Here *N* = 10. Note that planning using POMCP is intractable when *ε* = *−*3. (d,e) POMCP and ADP adaptively alternate between difficulty levels, thereby preventing catastrophic extinction. Note the drop in difficulty levels after significant extinction in both cases. Here *ε* = *−*2.

The POMCP teacher exhibits a non-monotonic curriculum, repeatedly reverting back to easier tasks before ramping up the difficulty. The *q* values for earlier steps in the sequence are relatively stable and lack the alternating reinforcement and extinction dynamics that we observe for the INC teacher (Figure 3d). This robustness extends to *ε* values lower than the critical value at which INC fails (*ε* =−2 in Figure 3c). Indeed, as shown in the example in Figure 3d, the POMCP teacher recognizes and compensates for significant extinction by rapidly decreasing the difficulty, increasing difficulty only after sufficient relearning occurs.

### D. A heuristic adaptive algorithm achieves near-optimal curriculum design

The POMCP teacher’s strategy suggests simple principles to overcome extinction while making learning progress. Specifically, a robust teacher algorithm has to 1) increase difficulty when the estimated success rate (*ŝ*) is sufficiently large (similar to INC), 2) continue at the same difficulty level when the success rate is below this threshold value as long as the student continues to learn (Δ*ŝ> µ*), and 3) decrease difficulty if the student begins to show signs of significant extinction (Δ*ŝ< µ*) for some *µ*. These three principles motivate our choice of a decision-tree-based teacher algorithm that uses *ŝ*and Δ*ŝ*as features. The precise splits and leaves of the trees can be optimized using various search procedures. More complex trees can be constructed by taking into account second or higher-order differences of the success rate. For the sequence learning task, we find that the features (*ŝ*(*t*), Δ*ŝ*(*t*)) are adequate to produce a successful teacher, which we term Adaptive (ADP). We optimize the decision tree using differential evolution (Figure 3b, see Appendix B 3 for details). Note that this optimized ADP is used for all benchmarks below with no additional tuning.

The ADP teacher shows dynamics similar to POMCP, mitigating extinction waves by alternating between difficulty levels (Figure 3e). We benchmark ADP against INC, POMCP and four algorithms proposed by Matiisen et al [41] (Figure 3c). These latter four algorithms are based on the principle of maximizing *learning progress* [43]: a student should attempt the difficulty level at which they make the fastest progress (as measured by the slope of the learning curve on a particular task). The algorithms differ in how progress is measured and how tasks are sampled based on their relative progress.

ADP is competitive with POMCP (for the range of parameter values that POMCP can be feasibly evaluated) and significantly outperforms the other algorithms for small values of *ε*, which is the regime where curriculum design is non-trivial and baseline algorithms such as INC fail. Moreover, ADP is robust when the innate biases for not equal (Figure S1). Since our OCL framework is task-agnostic and model-agnostic, ADP can be directly applied to other tasks and artificial agents provided that sub-tasks are arranged on a discrete, monotonic difficulty scale.

### B. Performance of ADP on deep RL tasks with delayed rewards

To examine whether ADP can design curricula for complex behaviorally relevant tasks and learning models, we train deep RL agents to solve two navigation tasks with delayed rewards: odor-guided trail tracking and plume-source localization.

Dogs are routinely trained to track odor trails, and various heuristics have been developed by trainers to efficiently teach dogs [50]. In a successful trail tracking episode, the student begins with a random orientation from one end of the trail and receives a reward only when they get to the other end of the trail. Trails are long, meandering and broken so that the agent is highly unlikely to get to the end through random exploration and should thus learn a non-trivial strategy to actively follow the trail and receive reward.

The trail tracking paradigm (Figure 4a-d) provides a natural split of tasks onto a difficulty scale. We design a parametric generative model for trails where the parameters control the length, average curvature and brokenness of the trails (Appendix C 5). Samples of trails along tasks of increasing difficulty are shown in Figure 4b. We develop a deep RL framework for trail tracking, where the tracking student uses its sensorimotor history of sensed odor and self-motion to modulate their orientation in the subsequent step (see Appendix C 5 for full details). Sensorimotor history is encoded using a visuospatial, egocentric representation (Figure 4a), so that the student has a memory determined by the size of the visuospatial observation window. The student uses a convolutional neural network architecture which is trained using Proximal Policy Optimization (PPO) [51].

**FIG. 4:**
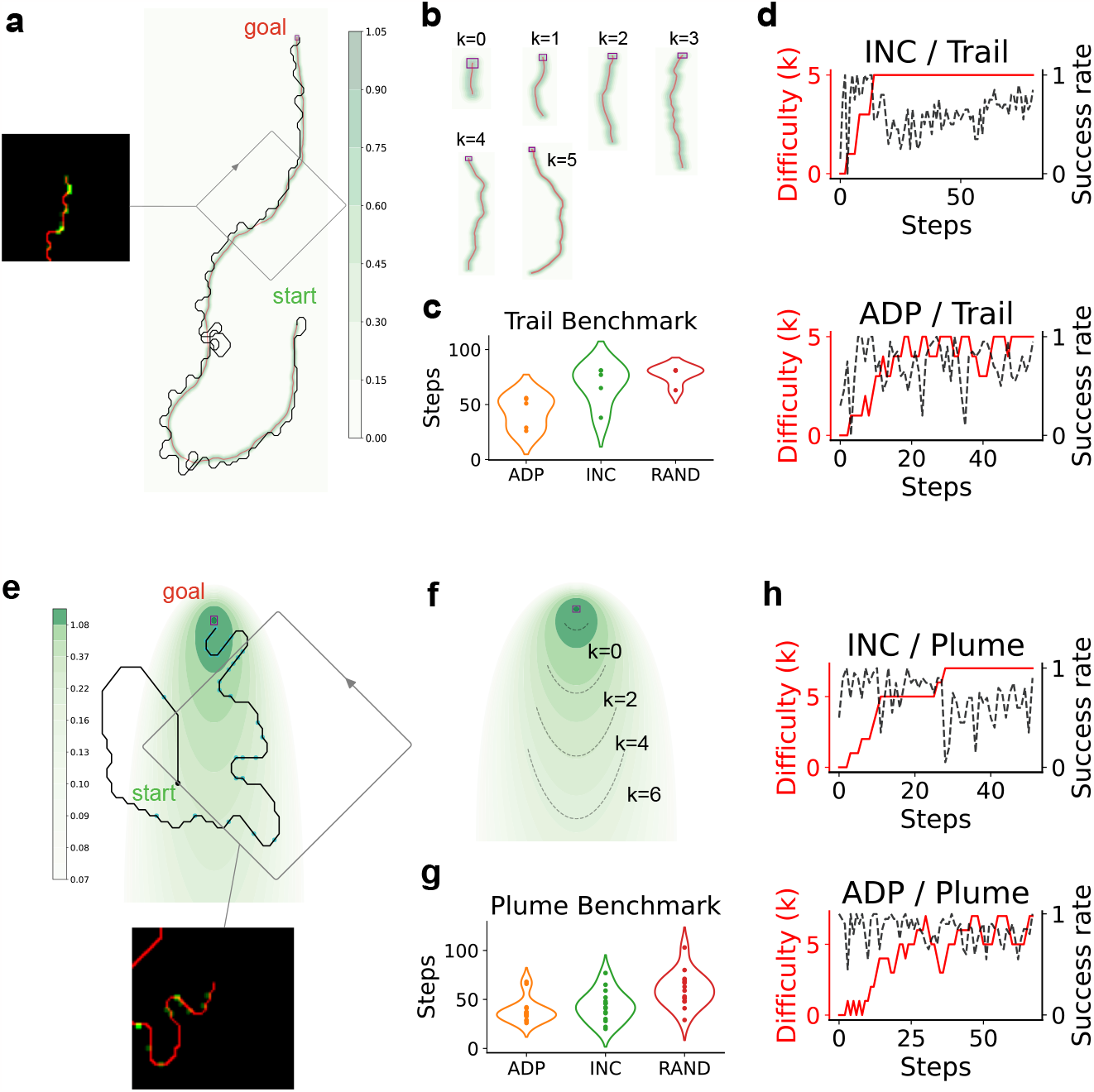
Deep reinforcement learning agents trained using a curriculum solve navigation tasks with delayed rewards. (a) The trail tracking paradigm. A sample trajectory of a trained agent navigating a randomly sampled odor trail. The colors show odor concentration. The inset shows the egocentric visuospatial input received by the network. (b) Sample trails from the six difficulty levels. (c)ADP outperforms INC and RAND (each teacher-student interaction is a step). Note that the agent does not learn the task without a curriculum. (d) The success rate of the agent in finding the target over training (black dashed line) for INC and ADP. The curriculum is shown in red. Note the significant forgetting shown by the student trained using INC approach compared to ADP. (e-g) As in panels a-d for a localization task. The agent is required to navigate towards a source which emits Poisson-distributed cues whose detection probability decreases with distance from the source (colored in green on a log scale).

The student does not learn without a curriculum. ADP outperforms both INC and a curriculum (RAND) that randomly chooses from the task set (Figure 4c). Their curricula show that ADP alternates as in the sequence learning task, presumably mitigating extinction effects associated with the transition to more difficult tasks. INC is comparable to ADP but experiences a greater degree of forgetting as seen in the longer time it spends at the highest difficulty level (Figure 4d). The path of an agent tracking the trail is shown in Figure 4a. The agent exhibits a preference for localizing at the edge of the gradient. When it encounters a break, the student performs repeated loops of increasing radius until it reestablishes contact with the trail. A detailed analysis of the student’s tracking behavior during trail tracking is postponed to future work.

Next, we extended this framework to a localization task (Figure 4e-g) inspired by naturalistic plume tracking [52, 53] and sound localization tasks. In each episode, the student begins at a random location a certain distance from a target whose (fixed) location is unknown. A unit reward is delivered when the student localizes at the target. The student receives sparse, Poisson-distributed cues from the target with probability that depends on the relative location to the target. These cues provide information about the location of the target, which can be used by the student to solve the task (see Appendix C 6 for full details). The delayed reward and sparse cues provide a challenge for training agents without a shaping protocol. We consider a curriculum where the difficulty scale is determined by the rate of detecting a cue at the student’s initial position, as well as the student’s distance from the target (Figure 4f). Similar to the trail tracking setting, results recapitulate the better performance of ADP compared to INC and RAND (Figure 4g,h).

### F. Continuous curricula

Our analysis up to this point assumes discrete curricula. A consequence of discrete curricula is that an unexpectedly large jump in difficulty from one level to the next can stall learning. In such situations, an animal trainer has the option of decomposing the task further and proceed with an INC approach. However, if the jump from one level to the next is too small, the student will progress in small steps while the teacher incurs a temporal cost on unnecessary evaluations. On a continuous curriculum, an optimal teacher has to adjust difficulty increments such that they reflect the student’s innate biases. Here, we explore preliminary ideas for designing continuous curricula using a continuous extension of the sequence learning task and a concomitant modification of the student’s learning algorithm (see Appendix D for more details).

We consider a continuous ADP teacher modified to accommodate the particulars of a continuous curriculum. At the start of an interaction, the experimenter proposes an initial “rough guess” for the difficulty increment used by the teacher. As the ADP teacher progresses, it tweaks the size of this increment based on the student’s performance. In addition to the three actions (increase, decrease and maintain difficulty), we introduce a second set of three actions: increase, decrease and retain the increment interval (the teacher selects from nine actions at each step). As in the discrete case, we use differential evolution to find the best decision tree (Figure 5a). Figure 5c,d shows trajectories for the continuous ADP teacher, which compares favorably with INC in benchmarks (Figure 5b).

**FIG. 5:**
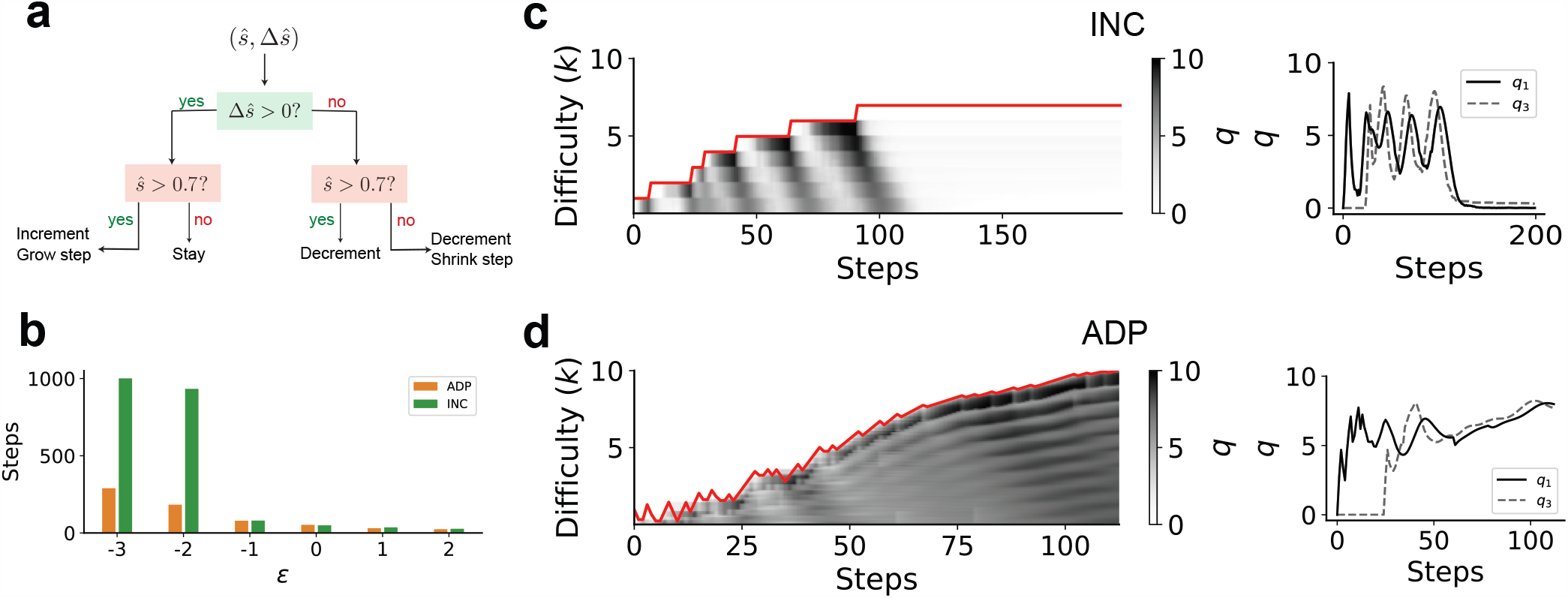
Algorithms for designing continuous curricula (a) Decision tree showing the continuous version of ADP which includes actions that “grow” and “shrink” the increments between continuously parameterized difficulty levels. See the text for more details of the task in the continuous setting. (b) ADP significantly outperforms INC when the task is difficult (low *ε*). (c,d) The *q* values plotted as in Figure 3d,e. Similar to the discrete setting, INC shows catastrophic extinction and never learns the task for sufficiently small *ε*. Continuous ADP first decreases increment size and smoothly increases the difficulty level while balancing reinforcement and extinction.

## IV. DISCUSSION

From Skinner’s missile guidance pigeons [54] to laboratory rodent experiments to state of the art artificial RL agents, curriculum design plays a foundational role in training agents to solve complex tasks. Here, inspired by behavioral shaping, we propose an outcome-based curriculum learning (OCL) framework and develop adaptive algorithms aimed primarily for training laboratory animals. In a sequence learning task, dual waves of reinforcement and extinction modulate the student’s performance, necessitating a careful shaping strategy that balances reinforcement and extinction. A naïve teacher, INC, fails to prevent extinction when students encounter large jumps in difficulty. A near-optimal teacher strategy (POMCP), discovered by formulating teaching as optimal planning under uncertainty, relies on frequent alternations between the current and previous task difficulty levels, which ameliorates extinction. Inspired by this observation, we use differential evolution to design a decision-tree-based heuristic algorithm, ADP. ADP is much more efficient and achieves performance comparable to that of POMCP, significantly outperforms other algorithms on the sequence learning task and requires no fine-tuning for the task or student. ADP outperforms other curriculum strategies when applied to train deep RL agents on complex, naturalistic navigational tasks.

We focus primarily on cases where the curriculum can be decomposed into rigid, discrete difficulty levels. Realworld tasks can often be further broken down when student’s encounter a bottleneck. We explore one continuous generalization of ADP that relies on finite approximations to continuous intervals, coupled with a *K*-step TD learning rule. The continuous setting poses a distinct challenge: since the teacher *a priori* does not know whether the student can solve an incrementally harder version of the task, estimating this through a transcript takes additional samples and thus incurs a temporal cost. Infintesimal increases in difficulty are not optimal. On the other hand, large jumps in difficulty will stall learning. We expect competitive algorithms to appropriately balance these two factors; a more exhaustive exploration of continuous OCL algorithms will be considered in future work.

The curricula we explore here have all involved a single axis of difficulty. For many real-world tasks, there are multiple axes that must all be optimized simultaneously. For example, a tennis player has to learn and compose multiple elements – footwork, various racquet motions, tactics – in order to improve general playing skill. In the trail tracking setting, we have simplified all such factors (length, average curvature, brokenness of the trails) into a single difficulty scale, when ideally, the teacher should choose how to modulate the difficulty along each factor.

One avenue for future work is to generalize our teacher algorithms to settings where there are multiple independent skills that need to be learned to solve the full task.

Finally, shaping is a crucial aspect of training animals. Concepts like task difficulty levels, innate bias *ε*, and behavioral extinction have natural analogs in biological agents. The teacher algorithms developed in this work can be readily deployed on real animals, and their efficacy measured. Many laboratory tasks in model systems such as mice involve extensive training lasting for weeks or more [13]. It is often unclear whether this lengthy training is due to the innate difficulty in animals learning the tasks at hand, or inefficient curriculum design [15]. Developing better teacher algorithms for animal training may result in significant savings in time and cost to produce well-trained subjects. In addition to practical benefits for laboratory research, any demonstration of more rapid training of animals will also shed light on their capabilities and limits of learning. Gradual shaping we discuss here may also be related to gradual introduction of more intuitive coincidences that exploit an animal’s priors to allow more rapid learning [15]. Such hand-crafted shaping is common in laboratory experiments [9–13], but more precise quantitative descriptions of behavioral learning algorithms such as ours opens the possibility of designing near-optimal teaching strategies in more general scenarios, similar to the POMCP formulation that we have developed here for a RL-based student. Optimistically, such formulations might even impact curriculum design for human students.

## ACKNOWLEDGMENTS

We thank Jacob Zavatone-Veth and members of the Murthy lab for helpful discussions. This work was supported by a joint research agreement between NTT Research Inc. and Harvard University, including grant A47994. VNM is partially supported by NIH RF1NS128865 and R01DC017311.

## Appendix A: Sequence learning

We describe in detail the specifics of the sequence learning task, and our model of a student. The task consists of a series of discrete choices, inspired by the setup in [44]. At each timestep, the student must select the correct action to advance. Selecting the wrong action at any timestep terminates the episode without reward. If the student advances *N* times in a row, where *N* is the pre-determined length of the environment, the episode terminates with fixed reward *R*.

Selecting the correct action is determined by the student’s *Q*-values, which are updated using a simple TD learning rule described below (Section A 1). Though the model is constructed with binary actions (correct / incorrect), it describes any situation where actions can be partitioned into correct and incorrect groups. The probability of selecting a correct action is modified using an additional parameter *ε*, which we describe further below. In this way, the task models any setting where a student must execute a consecutive sequence of correct actions, capturing a large portion of tasks classically suitable for curriculum learning.

See Figure 2a for a graphical depiction of the task. An MDP that summarizes this task can be given as

- **State space**: integers 1, 2, … *N*
- **Action space**: move forward, halt
- **Transitions**: For states *s, s*^*′*^ and action *a*, we have the following transition probabilities:

Pr(*s*^*′*^ = *i* + 1 | *s* = *i, a* = move forward) = 1, for *i < N*

Pr(*s*^*′*^ = ∅ | *s* = *N, a* = move forward) = 1

Pr(*s*^*′*^ = ∅ | *s, a* = halt) = 1, for any *s*

- **Reward**: the reward function is simply *R*(*s* = *N, a* = move forward) = *R* and 0 otherwise. For all sequence learning tasks, reward is fixed at *R* = 10. Note, for sufficiently large *R*, the specific choice of *R* will not qualitatively change the learning dynamics of the student. The student will learn quicker, but the learning curves will have the same shape. If *R* is chosen too small, the student experiences a bottleneck and its *Q*-values never saturate.

This setup lends itself naturally to a curriculum, where larger values of *N* correspond with more difficult tasks. The goal of the teacher is to propose a number of intermediate tasks indexed by their lengths *n*_1_, *n*_2_, *n*_3_, … such that the student succeeds at the final task of length *N* .

Note, we assume here that the task has already been broken down into *N* discrete steps. For a generalization to the continuous setting where *N* can vary along the real numbers, see Appendix D.

### Model of the student

A student in the sequence learning setting is modeled as an Expected SARSA RL agent [55, 56]. For each state *i*, the student has two *Q*-values: 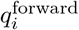 and 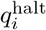, which correspond to the correct and incorrect actions, respectively. Because the incorrect halt action is never reinforced, we always have that 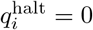. For simplicity, we omit specifying 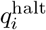 explicitly, and refer to 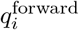 as simply *q*_*i*_.

In a standard softmax policy, the probability of moving forward from state *i* is given by

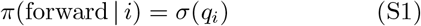

where *σ* is the sigmoid function, 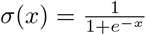 .

In the real world, different students have different propensities for learning a task. One lab mouse may be easier to train than another. One dog may have a better sense of smell, and track a trail more efficiently than another. One student may learn quicker than another, and follow a correspondingly accelerated curriculum. Modeling a student’s propensity for a task is therefore an essential consideration when designing curricula.

In our sequence learning setting, we incorporate this property by introducing a bias term *ε*. This term serves two related purposes:

1. **Model the student’s innate propensity for success**: every student has a different set of innate talents and preferences. Modulating *ε* allows us to simulate students with abilities that are more or less aligned with the task.
2. **Model the student’s extrinsic chance of success**: for more complex tasks, a student often has to choose one correct option among many incorrect choices. Modulating *ε* controls the student’s initial probability of success, capturing extrinsic factors that may aid or hinder the student’s progress.

To model these effects, we adjust the student’s policy to be

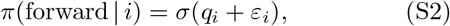

where *q*_*i*_ is updated through learning, but *ε*_*i*_ remains fixed. The value of *ε*_*i*_ can be different for different steps *i*, but in our experiments we typically set *ε*_*i*_ to be the same across all *i* and simply write it as *ε*.

The value of *q*_*i*_ is updated using the Expected SARSA update rule. Upon taking the correct action from state *i*, the value of *q*_*i*_ becomes

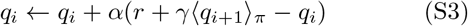

Note, the expectation ⟨ *q*_*i*+1 *π*_ ⟩is computed including bias *ε*, so

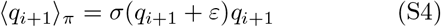

To simplify the analysis, we assume the student has an infinite horizon, i.e., the discount factor *γ* = 1.

## Appendix B: Teacher algorithms

We describe in greater detail our three primary teacher algorithms: Incremental (INC), POMCP, and Adaptive (ADP). Specific hyperparameter settings for all algorithms are attached at the end of this supplement.

### 1. Incremental teacher

An intuitive strategy for building a curriculum is to assume an incremental approach: the teacher proposes tasks of incrementally increasing difficulty as the student masters each successive level. For the sequence learning task, if the goal is to learn a task of length *N*, the teacher would start with a task of length 1. Once the student masters length 1, the teacher proposes length 2. Once the student masters length 2, the teacher proposes length 3. And so on until the student masters length *N* . See Algorithm 1 for a concrete description of this process. See Appendix E for details on how student’s are evaluated.

#### Algorithm 1 Incremental teacher for discrete curriculum

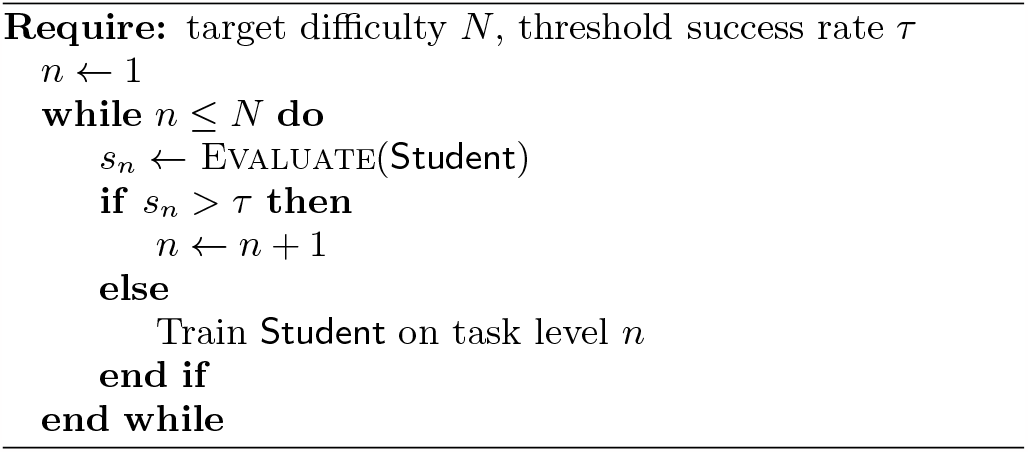

### 2. POMCP teacher

Just as the student is a reinforcement learning agent, the teacher can also be considered a reinforcement learning agent attempting to discover an optimal policy for a partially-observable Markov decision process (POMDP) [57]. Informally, the teacher’s observation space consists of the student’s outcome history up to the current time. After observing the student perform for a time, the teacher must take an action: namely selecting the next level of task to present. The process is partially observable because the teacher lacks full knowledge of the student’s state, and must infer the student’s true level of performance from the outcome history. Discovering an optimal policy for the resulting POMDP would shed insight on how an optimal curriculum should look for any particular student.

Formally, we define the following POMDP underlying the teacher’s decision process:

- **State space**: states consist of a 3-tuple (*q, ε, α*) where *q* represents all of the student’s learnable *Q*-values, *ε* represents the student’s innate ability bias, and *α* represents the student’s learning rate. In effect, the state is the set of values that fully define the student’s ability to perform the sequence learning task.
- **Action space**: integers 1, 2, … *N*, which correspond to the next task level that the student will see. To ensure that solving the POMDP remains tractable, we limit the action space to 3 values: 1) increment the task difficulty, 2) decrement the task difficulty, 3) keep the same task difficulty
- **Transitions**: transitions from (*q, ε, α*) to (*q*_new_, *ε, α*) can be sampled given a transcript by applying the SARSA update rule in Equation S3. For POMCP, we do not need an explicit probability distribution across transitions, only the ability to sample from it.
- **Reward function**: for a pre-determined success threshold *τ*, if a student’s rate of success at the final task exceeds *τ*, the teacher receives a fixed unit reward *R*. Concretely, the student’s rate of success at a sequence of length *k* is

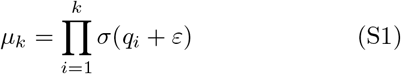

Let *s*^*\**^ = (*q*^*\**^, *ε, α*) be a 3-tuple for which *µ*_*N*_ *> τ* . Then any transition that terminates in *s*^*\**^ receives reward *R*, and the episode terminates. No other transition is rewarded, or terminates the episode (unless a pre-determined max number of iterations is exceeded).

- **Observation space**: the teacher observes the student’s transcript **h** = (*h*_*t*_, *h*_*t−*1_, …, *h*_*t−T* +1_) since the last interaction, where *h*_*i*_ = 1 if the student succeeded, and 0 otherwise. Because the transcript prior to the last interaction is encoded in the estimated parameters (*q, ε, α*), we do not need the entire history back to *t* = 0, and can keep just the last *T* episodes.

Further implementation details for POMCP are in Appendix F.

We use the Partially Observable Monte-Carlo Planning (POMCP) algorithm [48] to approximately solve this POMDP. The resulting teacher proposes curricula that differ from INC in several striking ways (Figure 3). Most importantly, rather than monotonically increment the curriculum by steady intervals, the POMCP teacher backtracks to earlier levels in an oscillatory movement. By alternating between two or more levels, the POMCP teacher pre-empts an extinction wave before it propagates all the way back to the start, erasing the student’s progress. In this way, the teacher encourages new reinforcement waves to form at higher levels while mitigating the impact of extinction, and successfully trains a student even for severely low *ε*.

### 3. Adaptive teacher

A key insight from the success of the POMCP teacher is that “backtracking” is essential to the success of a curriculum. Rather than increment in gradual, monotonic steps as the student learns, it is important to interleave easier levels periodically so as to counter extinction effects. Whereas POMCP decides the the points at which to backtrack through a blackbox search procedure, the Adaptive teacher explicitly optimizes for points to backtrack.

ADP works by learning a decision tree that decides whether to increment or decrement the current task level. As input, the teacher receives a history of success rates *s*^(1)^, *s*^(2)^, … *s*^(*t*)^ estimated from the student at each step up to the current time *t*. A *first-order* Adaptive teacher will additionally compute Δ*s*^(1)^, Δ*s*^(2)^, … Δ*s*^(*t*)^ where Δ*s*^(*i*)^ = *s*^(*i*)^ −*s*^(*i−*1)^ (and *s*^(*j*)^ = 0 for *j <* 1). When making a decision, the Adaptive teacher then compares the current success measures (*s*^(*t*)^, Δ*s*^(*t*)^) against a decision tree whose leaf nodes correspond to one of three possible actions: 1) increment task level, 2) decrement task level, and 3) stay at current task level. See Figure 3 for an example of what this decision tree looks like.

The precise splits and leaves of the trees can be optimized using any number of popular search procedures [58–61]. The Adaptive teacher can be further customized with additional “features.” For example, we could also implement a *second-order* teacher, which includes features Δ^2^*s*^(*i*)^ = Δ*s*^(*i*)^ − Δ*s*^(*i−*1)^, or arbitrary features Φ(*s*^(*i*)^) = *f* (*s*^(1)^, *s*^(2)^, … *s*^(*i*)^) for some arbitrary function *f* . For this simple sequence-learning task, we find that the features (*s*^(*t*)^, Δ*s*^(*t*)^) are adequate to produce a successful teacher.

Optimizing the Adaptive teacher proceeds as a coordinated ascent. The procedure begins with an initial, reasonable set of actions selected by the experimenter. Differential evolution [58] is used to evolve the precise splits in the tree, followed by a comprehensive search through the entire space of possible actions. These two steps, evolution followed by action search, alternate until converging on a final tree.

## Appendix C: Odor tracking

During olfactory navigation, an animal encounters a series of discrete odor measurements in pursuit of a target. Odors can be terrestrial or airborne, that is, concentrated along a single thin trail or diffused along turbulent air currents, respectively. Examples include a dog tracking the scent of deer, a moth following pheromones to find a mate, or a mouse pursuing fragrant hints of last night’s leftovers. This behavior is prevalent and essential across species, necessary for navigation, foraging, and mating [62–64].

Below, we detail two related deep reinforcement learning tasks that simulate odor-guided navigation: 1) surface-borne odor trail tracking and 2) airborne odor plume tracking. For both settings, we employ the same deep RL framework to recreate naturalistic tracking environments.

### 1. RL framework

Given the successes of various deep RL algorithms on video game tasks [65], we encode sensory-motor history using a visuospatial pixel representation. This representation allows the agent to learn both a sensory stack for interpreting the image inputs as well as a navigational policy for finding the target.

#### a. Observation space

For both trail and plume tracking, the agent is centered on a flat, 2D surface without landmarks or obstructions. Odor is either distributed along a thin trail (for trail tracking) or diffused in simulated plume (for plume tracking). As the agent navigates this landscape, it encodes its position history together with the last few odor detections within each individual image observation.

The observation space consists of a 50×50 grid of pixels and 3 color channels per pixel, which produces a full RGB image. Although the agent’s position is recorded continuously during each episode, when producing the pixel observations, it is discretized to the pixel grid. The three color channels of the image encode different information. The first channel (i.e. the “red” channel) encodes the agent’s position history. The second channel (i.e. the “green” channel) encodes the agent’s odor measurements. The last channel (i.e. the “blue” channel) is left unused.

At each timestep, the agent moves to a different location and performs an odor measurement. Position history is recorded as a thin continuous line in the image observation’s red channel, linearly interpolating between discontinuous points. Odor history remains discrete, and is depicted as colored patches at each location where the agent performed an odor measurement. This difference reflects the fact that motion through a space is continuous, but odor measurements occur only when the agent “sniffs” the environment. Hence, odor measurements in history must be discrete. The strength of the odor is proportional to the pixel value in the green channel. The image overall remains centered on the agent, that is, it is egocentric with respect to the agent’s position and heading. As the agent moves in a particular direction, the whole “viewport” moves and rotates with the agent, leaving a track that spreads out away from the center recording the agent’s past. See Figure 4a for an example observation that the agent receives.

**FIG. S1:**
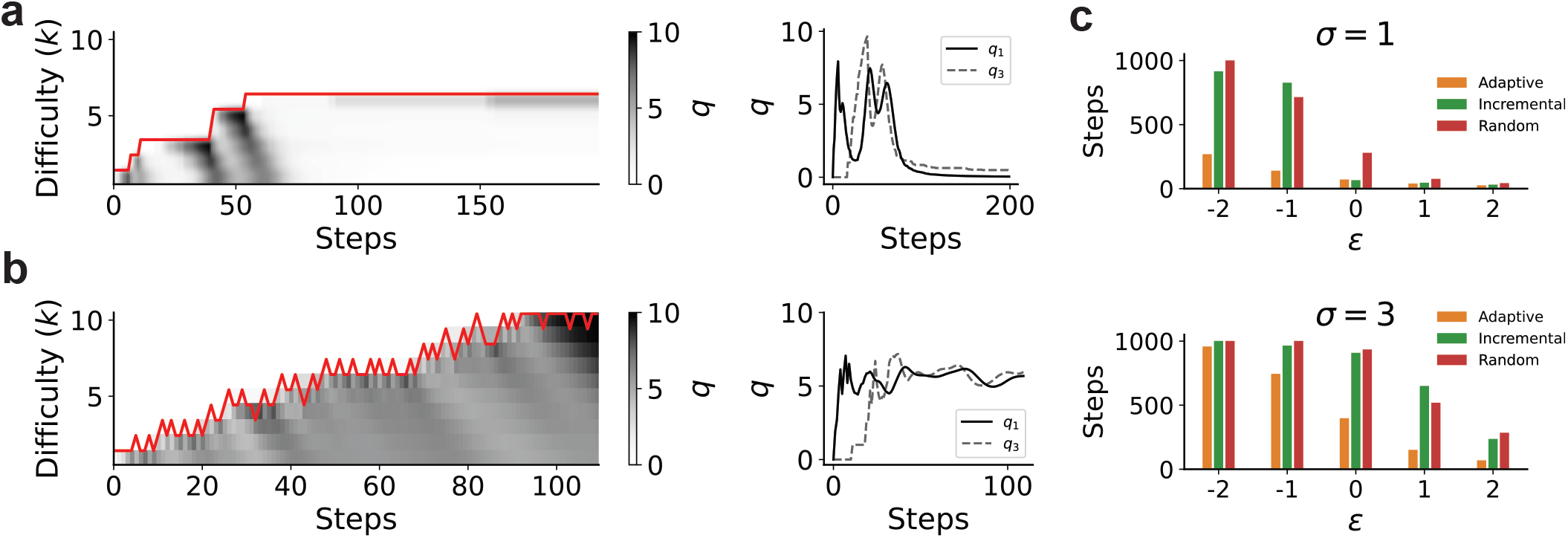
(a,b) *q* value dynamics for INC (panel a) and ADP (panel b) when *ε*_*i*_ values for each difficulty level are heterogeneous. Specifically, *ε*_*i*_ = *ε* + *ση*, where *η* is a standard normal random variable. These examples are run with *ε* = *−*1 and *σ* = 1. (c) Performance comparison between ADP, INC and a random strategy for *σ* = 1 and *σ* = 3.

### 2. Action space

At each timestep, the agent can take one of three actions: 1) move left, 2) move right, 3) move forward. Choosing the move left (or right) action will shift the agent’s heading by 45° in the corresponding direction. After any heading updates, the agent’s position is then incremented by three units along its new direction.

### 3. Reward

If the agent lands within 3 units of the target (either the end of a trail or source of a plume), the episode terminates with a large, fixed reward. Otherwise, if the agent fails to reach the target within a predetermined maximum number of iterations, the episode terminates without reward. No other actions are reinforced, nor are any negative rewards ever applied.

Note, the reward scheme is intentionally sparse. If an agent were to attempt a long trail without additional aid, it will fail to converge towards a successful strategy due to the lack of sufficient reinforcement signals. A traditional approach to addressing this sparsity is through reward shaping [66, 67], a process whereby the designer supplies supplementary rewards that guides the agent towards successful behavior. However, reward shaping is difficult to implement in practice, may require significant assumptions about the student, and different shaping strategies may have unpredictable impacts on the agent’s overall behavior [55, 68]. Instead, we use curriculum learning to overcome the sparsity issue. By starting the agent on short, easy trails for which the reward scheme is not sparse, then gradually lengthening the distance to the target, the agent naturally learns a successful tracking strategy without the heavy-handed tuning required for reward shaping.

### 4. Model

We use Proximal Policy Optimization (PPO) [69, 70], a popular deep RL algorithm that achieves state of the art on a wide variety of discrete and continuous tasks. The agent uses a deep convolutional neural network to extract operable features from each image observation, followed by several fully-connected layers to infer state values and output actions. For specific implementation details and hyperparameter settings, please see Appendix F.

### 5. Trail tracking

Surface-bound odor trails are long, thin segments of concentrated odor with minimal diffusion through the air. To sense these trails, an animal must be relatively close, and may not always sustain contact. Further, these trails often have breaks: significant stretches during which odor is absent, and the animal must execute a search strategy to regain contact. Terrestrial trail-tracking is therefore a highly nontrivial behavior.

To construct naturalistic trail geometries, we use the procedure described in Reddy *et al*. [71]. Trail characteristics are modulated with the following parameters:

- **Length**: the distance between the agent’s starting position and the target.
- **Width**: the scale at which odor diminishes along the axis perpendicular to the trail. With width parameter *σ*^2^ and a distance *x* from the trail, the concentration of odor *o* is proportional to

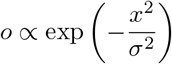

- **Heading**: the angle between the agent’s initial heading, and the direction of the target.
- **Shape**: the shape of the trail is governed by two parameters: curvature and diffusion rate. Curvature governs how the high-level shape of the trail evolves over time. Diffusion rate governs how “kinky” the trail is on short intervals.
- **Breaks**: a trail can have one or more breaks of variable length, along which there is no odor.

The agent perceives the exact magnitude of odor at its current location, which is represented in the magnitude of the green color channel in each image observation. This is in contrast to Reddy *et al*. [71], which uses a Poisson-based odor detection model. See Figure 4a for a plot of an example trail, including the trajectory of a successful agent.

For the purpose of building curricula, the difficulty of a trail is rated by a combination of its length, width, and breaks. The heading of the trail is allowed to vary across the entire compass rose, and the shape parameters are held constant across all episodes. Curricula are discretized by hand, with predetermined difficulty levels that specify particular parameter combinations. The specific difficulty levels are described in Appendix F. Because our teachers rely on discrete curricula, this extra manual intervention is necessary to apply these algorithms. In an ideal case, teacher algorithms should generalize to continuous curricula, which we explore further in Appendix D. At the start of every episode, a new trail is sampled using the parameter combination specified by the curriculum, and the agent must track an unseen trail from the beginning.

### 6. Plume-source localization

Often times, odor diffuses through the air in turbulent plumes. For humans, this mode of olfaction is perhaps more commonly experienced than terrestrial odor trails: the smell of baked goods at the local pastry shop, fragrance from a spring garden, or the comforting scent of your home are all plume-based odors. Like terrestrial trails, odor plumes are essential for survival and reproduction across animal species. However, tracking the source of an odor plume presents its own unique difficulties. Plumes tend to be rarefied and clumpy, where air turbulence partitions regions of odor concentration into random, disconnected patches. An animal attempting to locate the source of a plume must infer its location based on sporadic contacts and indirect cues.

To simulate naturalistic plumes, we use the model described in Vergassola *et al*. [72]. Odor plumes are shaped through the following parameters:

- **Wind speed**: a high wind speed produces long, elongated plumes. A low wind speed produces squat, round plumes. Zero wind produces a spherical plume.
- **Start rate**: the initial rate of detections at the agent’s starting position. A low start rate implies that the agent will start further away from the source of the plume, and have a correspondingly harder task.
- **Particle properties**: additional parameters like diffusivity, lifetime, and emission rate are related to the properties of individual particles, and influence the overall shape of the plume.

In contrast to the terrestrial trails, where the agent deterministically perceives the exact magnitude of odor at its current location, the odor detection model in the plume setting is probabilistic and discrete, accounting for the random influence of turbulence. Odor detection is Poisson-distributed, with the rate given by the plume model. See Figure 4e for an example plume, including the trajectory of a successful agent.

For the purpose of building curricula, the difficulty of a plume is rated by its start rate, with lower start rates corresponding to more difficult trails. To ensure the agent is always downwind of the source, the agent’s location is initialized within a fixed sector of a particular start rate. As in the terrestrial trail setting, because our teacher algorithms can only accommodate discrete curricula, we discretize the start rate into discrete levels for curriculum learning, with the specific levels given in Appendix F. A framework for handling continuous curricula is explored further in Appendix D. The agent’s starting *location* is re-sampled at the start of every new episode, though the starting *rate* remains the same for a particular task level. Particle properties and wind speed remain fixed for the entire duration of training.

## Appendix D: Continuous curricula

Introducing continuous curricula is a complex and nuanced topic. In this Appendix, we introduce a set of preliminary ideas for addressing the issue, but defer a more complete study to a future paper.

### 1. Continuous sequence learning

It is not immediately obvious how to generalize the sequence learning task to a continuous curriculum. Here, we propose one approximate scheme that preserves many (but not all) of the intuitions from discrete sequence learning, though it is by no means the only possible scheme.

We begin with the following notion. In the discrete sequence setting, for an untrained student with fixed bias *ε*, the probability that the student advances from step *n* to step *n* + 1 is given by *π*(*n* + 1|*n*) = *σ*(*ε*). Suppose the student is allowed to move continuously. Then the probability that the student first moves to 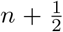, then from 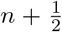 to *n* + 1 must also be *σ*(*ε*) overall. Hence, we must maintain that *π*(*n* + 1*/*2|*n*)·*π*(*n* + 1|*n* + 1*/*2) = *σ*(*ε*). In other words, there exists a value *ε*_1*/*2_ such that *π*(*n* + ½|*n*) · *π*(*n* + 1|*n* + 1*/*2) = *σ*(*ε*_1*/*2_)*σ*(*ε*_1*/*2_) = *σ*(*ε*).

In this example, we can see quite clearly that

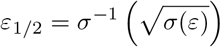

In the general case, if a student with *effective* bias *ε* moves an interval Δ*x* ≪ 1, then we can identify a value *ε*_Δ*x*_ such that *π*(*n* + Δ*x*|*n*) = *σ*(*ε*_Δ*x*_), where

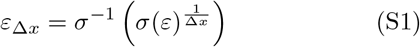

If we allow Δ*x*→ 0, then we approach a continuous curriculum. In the results that follow, we set Δ*x* = 0.01. In this way, we transfer much of the same intuition from the discrete sequence learning task. For an *effective* task length *N* and bias *ϵ*, which matches *N* and *ε* from a discrete curriculum, the student operates on a “continuous” environment with 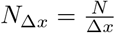 and *ε*_Δ*x*_. Ideally, a student operating on (*N*_Δ*x*_, *ε*_Δ*x*_) should perform identically to a student operating on (*N, ε*), as long as each curriculum increment is 1*/*Δ*x*. However, if we apply Equation S3 directly to update the student, we run into an important issue: only the previous *Q*-value is updated. For the continuous student, each *Q*-value represents an infinitesimal slice of ability. Applying the old rules as-is means we update only singular, infinitesimal slices at a time on the student. Intuitively, improving at a particular level should improve multiple *Q*-values simultaneously, and in a fashion that remains consistent with the discrete case. We model this effect by making a small change to the student’s update rule.

### 2. K-step student

For a student on our continuous approximation of the simple sequence learning task, the number of *Q*-values grows as the unit of discretization Δ*x* decreases. Indeed, for a curriculum of effective length *N*, the number of *Q*-values becomes 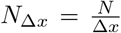. Without changes to the update rule described in Equation S3, the student is limited to updating one *Q*-value at a time. As Δ*x*→0, the number of *Q*-values to-be-updated approaches infinity, requiring infinite time for any reinforcement wave to propagate to the start.

Intuitively, as a student develops one particular *Q*value, neighboring *Q*-values should also be updated. After all, because *N* ≈*N*− Δ*x*, if *q*_*N*_ changes, *q*_*N−*Δ*x*_ should also change by approximately the same amount. Note, progress may be symmetric such that *q*_*N*+Δ*x*_ also change by a similar amount. Such an approach implies some form of a smoothing strategy for the update rule. However, for simplicity and consistency with the discrete case, we assume asymmetric updates where only *Q*-values less than the current task level *N* have an opportunity to be updated. A natural way to incorporate this intuition is to change from a single-step update rule as described previously to *K*-step expected SARSA.[55] That is, rather than update each *Q*-value using its immediate next neighbor, the update rule now becomes

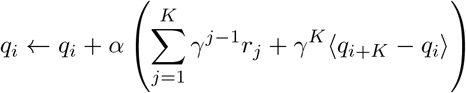

where *r*_*j*_ is the reward observed *j* steps ahead of the current step *i*. Because reward is only dispensed at the very end of a successful run, we have that *r*_*j*_ = 0 for *j < N*_Δ*x*_. Using a discount *γ* = 1 as before, the update simplifies to

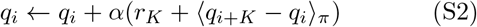

In effect, rather than updating from the immediate next-neighbor *Q*-value, the *K*-step student now updates using the *Q*-value *K* steps ahead. Hence, upon encountering a reward, all previous *K Q*-values are updated rather than just the immediately preceding one. On a later pass, as the student approaches this block of updated *Q*-values, the preceding block of *K* values also receives updates as the intervals overlap. If we allow *K* = 1*/*Δ*x*, the range of updated values correspond precisely with the updated values in the discrete sequence learning task.

For sufficiently high *ε*, the continuous *K*-step student corresponds precisely in learning speed with the discrete student. However, one important difference is the propagation of mistakes. In the discrete case, if the student halts, the error is not backpropagated. Rather, the (unwritten) *Q*-value associated with choosing the halt action is “updated,” and remains 0. However, in the *K*-step scenario, because all previous *K Q*-values are updated, if the student halts, the last *K*−1 *Q*-values are depressed slightly with a zero update, initiating a secondary extinction wave distinct from the ones generated through curriculum changes. Hence, learning is slower and more difficult for continuous students, and the potential for extinction waves to erase all progress correspondingly higher. The situation is exacerbated by low *ε*, in which the probability of halting is higher. An easy way to fix this situation is to simply alter the learning rules such that halting actions do not propagate depressed *Q*-values. However, it remains unclear whether this effect is a feature or bug — perhaps it is more realistic for repeated failures to depress the performance of the student, rather than leave performance unimpacted. Perhaps the correct action is to modify the discrete student such that errors also propagate. Ultimately, we leave this distinction as an additional layer of complexity for the continuous student, and an additional obstacle an optimal teacher must overcome.

### 3. Teacher algorithms

We explore generalizations of both the Incremental and Adaptive teachers. Unfortunately, POMCP does not scale to the massive size of the POMDP in the continuous case. We rely on the behavior of the Adaptive teacher to give a sense for what optimality looks like in this setting. Note, none of the algorithms from Matiisen *et al*.[41] generalize at all to continuous curricula, so we do not investigate them further in this setting.

Incremental generalizes directly to this setting. We allow the curriculum to increment in fixed intervals of 1*/*Δ*x*, monotonically increasing the task difficulty as the student progressively attains mastery. The procedure otherwise remains identical to the one described in Algorithm 1. Figure 5 shows trajectories for the Incremental teacher adapted for the continuous setting.

#### a. Continuous Adaptive teacher

The Adaptive teacher is similar in spirit to its discrete counterpart but extended to accommodate the particulars of a continuous curriculum. At the start of an interaction, the experimenter proposes an initial “rough guess” at an appropriate increment interval for the teacher to use. Such an increment can be considered a prior that the experimenter assumes about the difficulty of a task. For the continuous sequence task, we use an initial increment of 1*/*Δ*x*. As the Adaptive teacher progresses, it tweaks the size of the increment according to the student’s performance. Hence, rather than just three actions (increment, decrement, stay), we introduce a second set of three actions: increase the increment interval, decrease the increment interval, retain the same interval. Adjustments to the interval are made multiplicatively by a predetermined percentile.

With this adjustment, at every iteration, the Adaptive teacher must now select from one of nine possible actions: an `increment, decrement, stay`, paired with a `grow interval, shrink interval`, and `keep interval`. To select the correct action, one can use the same optimization procedure described in Section B 3 to learn the best actions for each situation. Figure 5 shows trajectories for the Adaptive teacher adapted for the continuous setting (compare to Figure 3e).

#### b. Benchmarks

See Figure 5 for a comparison of Incremental and Adaptive teachers on the continuous sequence learning task. Adaptive has a decisive edge over Incremental, particularly for high *N* and low *ε*. Random and finaltask-only curricula are not plotted, as learning fails to occur even for the easiest tasks pictured.

Overall, intuitions from the discrete case translate naturally to the continuous case, though learning overall tends to be more difficult in the latter. We presented here only an initial exploration of one particular continuous generalization. Future work will need to examine more deeply nuances particular to the continuous case, what optimal teachers may look like in this setting, and validate these results on a naturalistic task like trail tracking.

## Appendix E: Estimation methods

A central aspect of outcome-based curriculum learning is that the teacher does not have access to the student’s internal parameters. Rather, the teacher must estimate intrinsic qualities of the student through extrinsic observables alone. In our context, the critical internal parameter of the student we estimate is its true success rate *s*_*n*_ for a task of difficulty *n*. For example, in the case of the Incremental teacher, if *s*_*n*_ exceeds some threshold *τ*, the student advances. Otherwise, the student remains on the current task level. In the case of Adaptive, *s*_*n*_ and Δ*s*_*n*_ are used as inputs to a decision tree that decides the student’s next task.

In all these cases, the teacher has access only to the student’s transcript of successes and failures: **h** = (*h*_1_, *h*_2_, …, *h*_*t*_), where *h*_*i*_ = 1 if the student succeeded on round *i*, and 0 otherwise. Given this transcript, the teacher must construct an estimate *ŝ*_*n*_(**h**) ≈*s*_*n*_. In the main text, we consider only a simple exponential moving average (EMA) approach to computing *ŝ*_*n*_. In this appendix, we describe the EMA in further detail, and explore two alternative estimation procedures motivated by a Bayesian approach.

### 1. Overview

Under the student model described in Section A 1, the true success rate of the student on a task of length *n* is given by

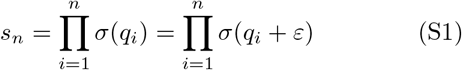

However, because the underlying parameters *q*_*i*_ and *ε* are unknown to the teacher, the quantity *s*_*n*_ must be estimated from the student’s transcript of successes and failures. Below, we compare three estimation procedures for *s*_*n*_:

#### 1. Exponential moving average

we apply an EMA to the student’s transcript to estimate *s*_*n*_. This estimate functions as a baseline with which to compare our other two, Bayesian motivated approaches

#### 2. Beta posterior inference

we assume the successes and failures in a student’s transcript are drawn iid from a Bernoulli distribution. Because the student’s success rate evolves as it learns, *s*_*n*_ is a nonstationary quantity, and so this assumption cannot hold over long periods of time. However, to simplify posterior inference, we assume that negligible training occurs over short time periods, and that *s*_*n*_ is locally stationary.

#### 3. Particle filtering

we relax the local stationarity assumption from Method 2 and develop a particle filtering algorithm to estimate *s*_*n*_. Particles are sampled from priors over *q*_*i*_ and *ε*, then forward-simulated to yield posterior samples over *s*_*n*_.

In the following sections, we develop each method in detail, and compare their ability to estimate *s*_*n*_. To remove the confounding influence from complex teacher strategies, we use only the Incremental teacher as a framework for comparing these estimation procedures, and later compare Incremental (with sophisticated estimation techniques) to the gold-standard POMCP teacher.

### 2. Exponential moving average

Recall that at time *t*, the teacher observes a transcript of the student’s performance **h** = (*h*_1_, *h*_2_, … *h*_*t*_), where *h*_*i*_ = 1 if the student succeeded on episode *i*, and 0 otherwise. The EMA algorithm constructs an estimate *ŝ*_*n*_ through the following recursive update rule as new observations *h*_*t*_ are made:

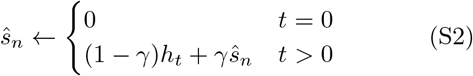

which corresponds to the unrolled equation

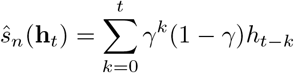

The parameter *γ*∈ [0, 1] is chosen by the experimenter prior to computing the EMA, and controls the degree to which past observations are discounted.

The exponential moving average is a simple, nonparametric approach to estimating a nonstationary quantity. See Figure S2a for a plot of EMA estimates compared to the true underlying success rate on a task with length *N* = 10, using the teacher model described above. We compare EMA across different discounts *γ* and across students with different bias parameters *ε*. Higher values of *γ* resulted in smoother estimates, though with a greater lag behind the true value. Lower values of *γ* resulted in noisier estimates, though with less lag. For *γ <* 0.5, the estimates became meaningless as they tended to alternate between extremes. Overall, a discount of *γ* = 0.8 seems to be the most appropriate, and tracks the true success rate with reasonable consistency.

### 3. Beta posterior inference

We next consider a simple Bayesian approach to this estimation problem. One challenge of estimating *s*_*n*_ is that this quantity is nonstationary. As the student learns the task, *s*_*n*_ changes over time. However, to make the analysis simpler, we can make the (big) assumption that over short intervals, *s*_*n*_ is essentially stationary. In this case, suppose *s*_*n*_ is approximately stationary over the time interval [*t*−*k, t*], for some small integer *k*. Then for a transcript **h** = (*h*_1_, *h*_2_, … *h*_*t*_), we have

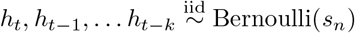

Hence, if we impose the prior *s*_*n*_ ∼ Beta(*α, β*), the posterior on *s*_*n*_ becomes

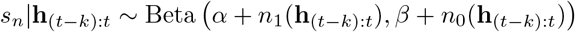

where *n*_1_ and *n*_0_ count the number of ones and zeros respectively. This form suggests an estimator based on the posterior mean 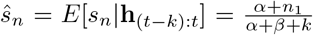 In this setting, we assume a uniform prior *s*_*n*_ ∼ Beta(1, 1). Se-\ lecting the value *k* is somewhat trickier. We would ideally like to select a *k* that is as small as possible, so as to ensure *s*_*n*_ does not change too much over the interval [*t*−*k, t*]. At the same time, if *k* is too small, our posterior has a higher variance, and the confidence in our estimate is correspondingly lower.

To address the latter issue, let us first consider the lowest value of *k* we can tolerate. Suppose we use a threshold *τ* such that if *s*_*n*_ *> τ*, then the student advances to the next level. Suppose we would like to be confident at the level *c* such that *s*_*n*_ exceeds *τ* before allowing the student to advance. Then we stipulate that *p*(*s*_*n*_ |**h**_(*t−k*):*t*_ *> τ* ) *> c*. In the best case scenario, all observations in **h** are ones. The minimum *k* we can tolerate is therefore the smallest *k* such that *p*(*s*_*n*_|**h**_(*t−k*):*t*_ *> τ* ) *> c* is true when **h** = 1, 1, … 1. To compute this *k*, we observe that

**FIG. S2:**
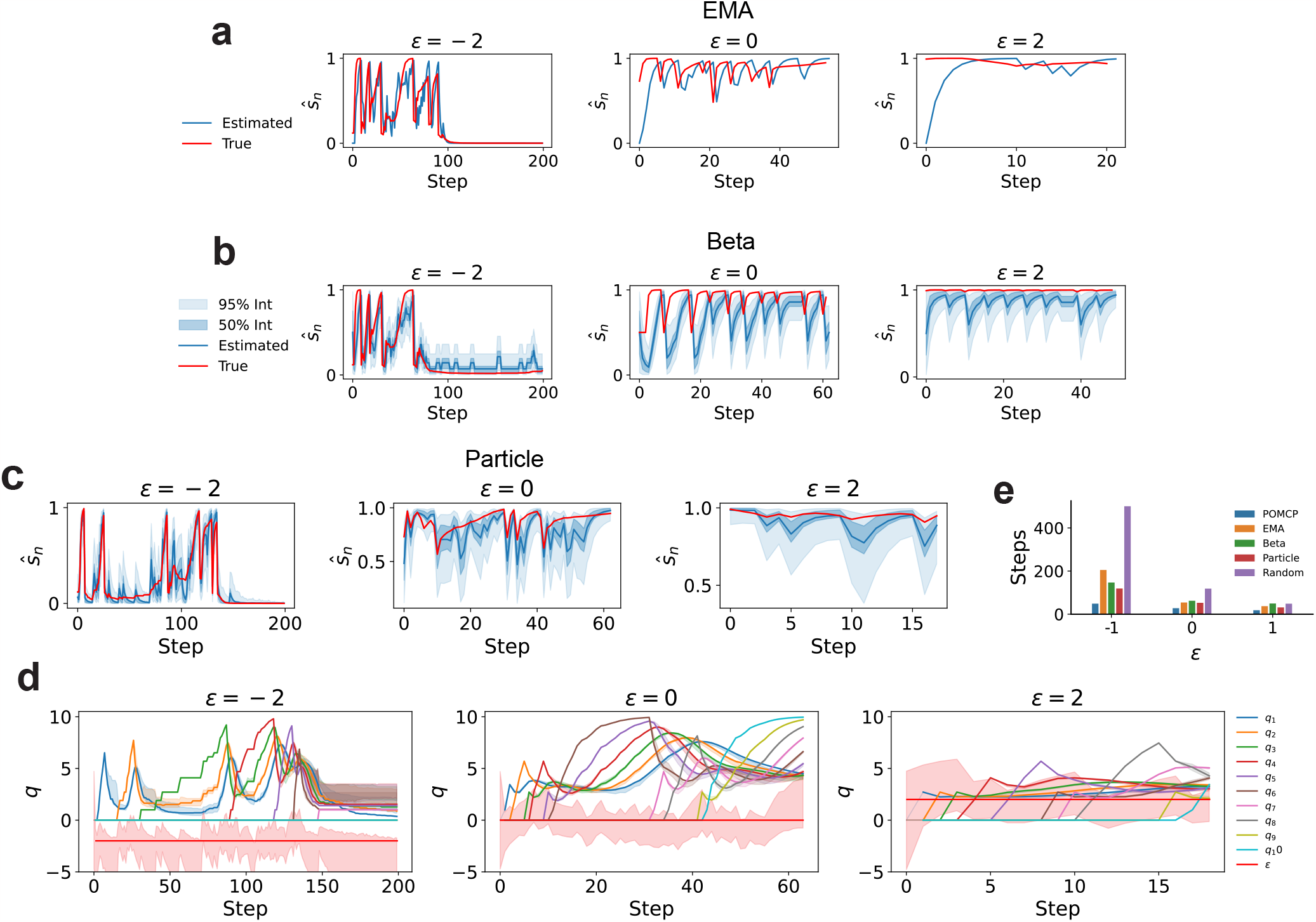
The following plots compare the performance of each estimation method under an INC teacher. (a,b,c) The true and estimated success rate when an exponential moving average (a) beta posterior inference (b) and a particle-filter-based approach (c) is used for estimation. In panels b,c the 50% and 95% confidence intervals are also shown. (d) The *q* values estimated by POMCP, with 95% intervals. (e) Performance comparison between ADP using the three different estimation methods. We use EMA in the main text due to its simplicity and efficiency.

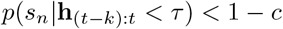

and

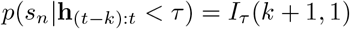

where *I*_*τ*_ (*α, β*) is the incomplete Beta function with upper limit *τ* . From here, we see that *I*_*τ*_ (*k* + 1, 1) = *τ*^*k*+1^ *<* 1 − *c*. Solving for *k* yields the final result

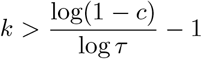

so we can establish a lower bound on *k* as 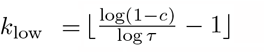. In practice, we use a confidence *c* = 0.5, which corresponds to when the median of the posterior exceeds the threshold *τ*, and yields reasonable results. Note, because *s*_*n*_ changes for different difficulty levels *n, k* should not be so large that it dips into episodes where *n* was smaller. Hence, after the student advances, we must wait at least *k*_low_ rounds before evaluating the student. For the EMA algorithm, this was a non-issue because EMA would adapt its estimate of the success rate to the current statistics. In contrast the Beta posterior procedure relies on an underlying stationary quantity, and so would be obviously invalid if it uses samples from *s*_*n−*1_ in its estimation of *s*_*n*_.

To determine what the upper bound on *k* should be, it is unclear how to select a reasonable upper-bound analytically. However, because *s*_*n*_ is presumably increasing over long timescales, the further back we look, the lower our estimate *ŝ*_*n*_ will become. Hence, we establish a coarse upper bound *k*_high_ = 3*k*_low_, and evaluate *p*(*s*_*n*_ |**h**_(*t−k*):*t*_ *> τ* ) *> c* for every *k* in between *k*_low_ and *k*_high_. If at any point this expression evaluates to true, we advance the student to the next level. Because *ŝ*_*n*_ decreases for larger values of *k*, there should ideally be a single *k* between *k*_low_ and *k*_high_ that balances a large enough sample that produces a high-confidence estimate, and a small enough sample that does not yield an underestimate on *s*_*n*_. As before, *k*_high_ should not be so large as to contaminate the data with samples from *s*_*n−*1_. The prefactor 3 was chosen as a hyperparameter setting that seems to work well in practice, though other factors may be used.

See Figure S2b for a plot of the Beta posterior estimate, along with the true estimates of the student’s performance. In general, the Beta posterior estimate tracks the trajectory of the student’s true success rate, but underestimates it systematically. This may be in large part because our assumption of local stationarity on the scale of *k* is invalid. Significant learning likely occurs even within just *k*_low_ steps, motivating the need to develop an approach that does not rely on this tenuous assumption.

### 4. Particle filtering

Our final estimation approach does not assume any stationarity in *s*_*n*_; rather, it estimates *s*_*n*_ by simulating the underlying learning dynamics using a particle filtering algorithm. Specifically, we estimate posteriors on the student’s *Q*-values *q*_*i*_ and innate bias parameter *ε*, then construct an estimate of *s*_*n*_ based on Equation S1.

The particle filtering algorithm proceeds as follows:

#### 1. Sample initial particles from the prior

We apply a uniform prior on *ε* over the interval [−5, 5], which encompasses the range of reasonable values *ε* can take. We apply a point-mass prior centered on 0 for all *q*_*i*_, because each *q*_*i*_ is initialized to 0 for every student. An initial set of particles 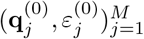 are sampled from these priors. In the simulations, we find that *M* = 1000 is sufficient.

#### 2. Simulate forward dynamics

For each particle, we simulate *T* trials of the student using the learning rules described in Equation S3, obtaining a set of updated parameters 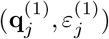 along with a set of corresponding simulated transcripts 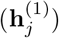

#### 3. Filter particles consistent with observation

After receiving an observed transcript **h** from the student, we keep all particles *j* such that 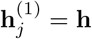. Note, contrary to previous descriptions where we assume **h** corresponds to the entire history of the student’s performance, here we assume **h** corresponds only to the student’s trials since the last interaction, and |**h**| = *T* . If *T* is small, checking direct equality works well. However if *T* is large, the probability that an observation will match a simulated transcript diminishes accordingly (even if the underlying parameters match), in which case we might compare a summary statistic on **h** like the mean. In our case, *T* = 3, which is sufficiently small to check for equality directly.

#### 4. Resample remaining particles

For our remaining particles, to remove any outliers, we perform a resampling weighted by each particle’s likelihood. For a particle 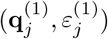 and observation **h**, if the student’s learning rate *α* is sufficiently small, the likelihood of the particle can be given as

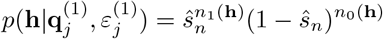

where *ŝ*_*n*_ is the particle filtering estimate of the true success rate at level *n*, and follows from equation S1

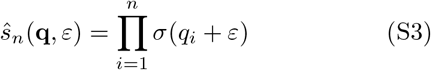

With these weights, all particles are resampled until we have a set of the original size *M* particles.

#### 5. Particle reinvigoration

Particularly for long runs, *ε* will tend to drift over time as unexpected observations are encountered (the *Q*-value parameters tend to remain fairly close, as we will see below). To forestall any posterior collapse, it is essential to reinvigorate *ε*. We use a simple reinvigoration strategy where a small random jitter is applied to every *ε* parameter. Specifically, we use a normally-distributed jitter centered at 0 with variance 0.25.

#### 6. Repeat

Encountering one observation yields a set of particles 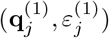, which represents samples from the posterior **q**, *ε*| **h**^(1)^. Upon encountering a second observation, we repeat our calculations from step 2 using the current posterior sample to obtain a new set of particles 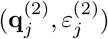, which represents samples from the posterior **q**, *ε* |**h**^(2)^, **h**^(1)^. This process is repeated for all new observations as they arrive.

Using this particle filtering algorithm, we generate posteriors on the parameters *q*_*i*_ and *ε*. At step *i*, the posterior samples 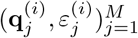 can be used to construct posterior samples on estimated success rate *ŝ*_*n*_ using Equation S3. From here, we can adopt the same approach as before, and if more than *c* proportion of the posterior samples are greater than a predetermined threshold *τ*, we advance the student to the next level. As for the Beta posterior inference approach, we use *c* = 0.5.

See Figure S2c for a plot of the estimated success rate during runs using the particle filtering algorithm. In general, the estimates fall quite close to the true success rate, outperforming both of the previous methods. Figure S2d plots 95 percent intervals on the posterior samples of the underlying parameters, compared to the true values for each parameter. The particle filtering approach tends to capture the true values well, confirming that its inference is accurate.

### 5. Comparison of estimation procedures

Figure S2e shows a comparison across all teacher algorithms, including the POMCP teacher. The POMCP teacher represents the approximately optimal teacher, and shows the lower bound that all other teachers approach. In general, our simple incremental teacher model approaches the optimal, though of course does not quite attain it, particularly for lower *ε* and higher difficulty levels *N* . Somewhat surprisingly, it also appears that the accuracy of an estimation method does not matter critically, as all three estimation methods produce students with more-or-less the same efficiency, though the particle-filtering method retains a slight edge.

In application, the particle filtering approach is no longer tractable as a success estimation method in more complex settings like our naturalistic tracking tasks. Between the remaining two options (EMA and Beta posterior inference), EMA is faster and simpler, with fewer tunable hyperparameters, while retaining good performance. Hence, we select EMA as our estimation method of choice in the main text.

## Appendix F: Implementation details

All code is available on GitHub: https://github.com/wtong98/automated-curriculum-learning

### 1. Sequence learning

In the sequence learning setting, the student is an expected SARSA agent with the following parameters:

| Parameter | Value |
| --- | --- |
| Reward | 10 |
| Learning rate | 0.1 |
| Discount | 1 |
| Episodes per interaction | 3 |

### 2. Success estimation

INC and ADP use an exponential moving average (EMA) to measure the success rate of the student, with discount factor *γ* = 0.8. (Alternative approaches are discussed in Appendix E.)

In the sequence learning setting, the EMA averages over the student’s history of successes and failures. In the deep RL setting, multiple students are run simultaneously to parallelize sample generation during rollout. The transcript across all parallel students are averaged in time to produce a single mean transcript for EMA.

### 3. Incremental teacher

The Incremental teacher has a single parameter: the success rate *τ* beyond which the student should advance to the next level. We use *τ* = 0.95 for the sequence learning task, and *τ* = 0.7 for the deep RL tasks.

### 4. POMCP teacher

The POMCP teacher uses the following parameter settings:

| Parameter | Value |
| --- | --- |
| Particles | 5000 |
| Discount | 0.9 - 0.95 |
| MCTS stop threshold | 0.01 |
| MCTS explore factor | 1 |
| Reinvigoration probability | 1 |
| Reinvigoration scale | 0.5 |

We implement POMCP using the particle filtering algorithm described in Appendix E, with an additional state variable *α* that represents the student’s learning rate. Particle reinvigoration applies only to the estimation of the student’s innate bias parameter *ε*. For a particle estimate *p*_*ϵ*_, reinvigoration proceeds as *p*_*ε*_←*p*_*ε*_ + *η*, where *η*∼ 𝒩 (0, *σ*). Estimation of the student’s learned Q-values tend to be highly accurate, and hence do not require reinvigoration.

A random rollout policy does not scale well to low *ε* for our problem. Instead, rollouts are computed following an “Incremental policy” – the decisions that an Incremental teacher would make. In other words, if the success rate of the student based on the sampled parameters exceeds *τ* = 0.95, the increment action is chosen. Otherwise, the stay action is chosen. An Incremental policy proved to work sufficiently well even for low *ε* tasks, though given the sensitivity of Incremental teachers to very low *ε*, a more intricate policy may perform more efficiently in this regime.

### 5. Adaptive teacher

Figure 3 shows the decision tree used by the Adaptive teacher on the sequence learning task (for both homogeneous and heterogeneous *ε*). The decision tree learned by the Adaptive teacher for the continuous case is shown in Figure 5. For the sequence learning task, optimizing the Adaptive teacher proceeds as a coordinated ascent. The procedure begins with an initial set of actions guessed by the experimenter. Differential evolution [58] is used to evolve the precise splits in the tree, followed by an exhaustive search through the space of possible actions. These two steps, evolution followed by action search, alternate until converging on a final tree.

The decision tree used by ADP on the deep RL tasks was tuned by hand, and proceeds as follows. For an estimated success rate *ŝ*and change in success rate Δ*ŝ*

- If *ŝ>* 0.7 and Δ*ŝ*≥ 0: increment
- If *ŝ<* 0.65 and Δ*ŝ<* 0: decrement
- Otherwise: stay

### 6. Matiisen teachers

We use a grid search to identify optimal hyperparameters for the teacher algorithms described in Matiisen *et al*. [41]. The final parameters we used are:

- **Online**: *α* = 0.05, *β* = 0.34
- **Naive**: *α* = 0.16, *β* = 3.8
- **Window**: *α* = 0.26, *β* = 4.83, *k* = 10
- **Sampling**: *α* = 0.1, *k* = 3

### 7. Trail tracking

Trails are generated using generalized worm-like chain ensembles, using the procedure described in Reddy *et al*. [71]. The parameters used to sample each trail are

| Parameter | Value |
| --- | --- |
| Width | 5 |
| Diffusion rate | 0.02 |
| Curvature radius | 70 |
| Heading | 0 - $2\pi$ |
| Break point | 0.5 - 0.6 |

The break point parameter specifies the segment of trail over which there is no odor, and is expressed as a proportion of the trail’s total length. The schedule used to produce Figure 4 varies based on the trail’s length. The specific lengths used are:( 10 30 50 70 90 100)

The agent is a PPO [69] deep reinforcement learning model with the following hyperparameters “MLP” refers to a multi-layer perceptron with 2 layers, 128 units per layer, and ReLU activations.

| Parameter | Value |
| --- | --- |
| Samples per rollout | 8192 |
| Batch size | 256 |
| Epochs | 5 |
| Learning rate | 1e-4 |
| Feature model | Nature CNN[73] |
| Action model | MLP |
| Value model | MLP |
| Entropy coefficient | 0.1 |
| Discount | 0.98 |
| GAE weight | 0.9 |
| Gradient clip range | 0.2 |
| Max gradient norm | 1 |
| Value function clip range | 0.36 |

The agent receives as input a pixel observation of the world with a square view distance of up to 40 units in the horizontal or vertical directions, scaled up by a factor of 2 to produce the direct pixel observations. At each step, the agent advances three units in the forward direction, or 45 degrees to the left or right. The agent’s heading rotates left/right by the same angle. The agent is allowed up to 200 steps before the episode terminates.

### 8. Plume tracking

Plumes are generated using the plume model in Vergassola *et al*. [72]. The parameters used to generate each plume are

| Parameter | Value |
| --- | --- |
| Length scale | 20 |
| Diffusivity | 1 |
| Emission rate | 1 |
| Particle lifetime | 150 |
| Wind speed | 5 |
| Sensor size | 1 |

The schedule used to produce Figure 4 varies based on the starting odor detection rate. The specific rates used are computed as 1*/r*_*k*_, where *r*_*k*_ = 0.5 + 0.1*k* and *k* increases from 0 to 23, for a total of 24 difficulty levels.

As before, the agent is a PPO [69] deep reinforcement learning model, with the following hyperparameters “MLP” refers to a multi-layer perceptron with 2 layers, 128 units per layer, and ReLU activations.

| Parameter | Value |
| --- | --- |
| Samples per rollout | 8192 |
| Batch size | 256 |
| Epochs | 5 |
| Learning rate | 1e-4 |
| Feature model | Nature CNN[73] |
| Action model | MLP |
| Value model | MLP |
| Entropy coefficient | 0.25 |
| Discount | 0.98 |
| GAE weight | 0.9 |
| Gradient clip range | 0.2 |
| Max gradient norm | 1 |
| Value function clip range | 0.36 |

The agent’s mechanics are identical to those of the trail case. The max steps the agent can take in the environment scales as 3 times the starting distance to the odor source.

## Notes

### Competing Interest Statement

The authors have declared no competing interest.

